# Alveolar Cytochrome P450 Mediates Butylated Hydroxytoluene-Induced Electrophilic Injury and Apoptosis

**DOI:** 10.64898/2026.09.08.750144

**Authors:** Michael T. Forrester, Katherine M. Harbaugh, Naoya Miyashita, Kyoshiro Tsuge, Rhianna Lee-Ferris, Jeremy M. Morowitz, Aleksandra Tata, Robert M. Tighe, Matthew W. Foster, Christine E. Eyler, Purushothama Rao Tata

## Abstract

Butylated hydroxytoluene (BHT) is a synthetic phenolic antioxidant utilized as a preservative in many products from foods to cosmetics. Population-based studies have detected BHT in the vast majority (>90%) of human specimens (serum, urine, fingernails) as well as home dust samples. Thus, BHT constitutes a ubiquitous environmental contaminant without obvious adverse health effects to humans. In contrast, exposure of mice to a single intraperitoneal dose of BHT triggers distal epithelial damage including loss of gas-exchanging alveolar type 1 epithelial cells, thus providing an invaluable tool to study alveolar repair and transient fibrosis. Presently, the molecular basis of BHT lung toxicity remains unknown. To address this, mouse lung single cell transcriptomic data were used to identify cytochrome P450 2B10 (CYP2B10) in AT1 cells as a BHT-activating enzyme. In cell culture systems, expression of CYP2B10 leads to marked BHT sensitization consistent with bioactivation of BHT into a toxic quinone methide. Targeted and proteome-wide experiments identify BHT-induced protein alkylation, DNA damage response, stress kinase activation and intrinsic apoptosis in a CYP2B10-dependent manner. Structure-activity relationship studies reveal the necessity of *para*-methyl and *ortho*-*t*-butyl groups necessary for BHT bioactivation. Our findings elucidate the molecular basis by which BHT is converted from an innocuous antioxidant into a highly toxic quinone methide and identify the first mammalian enzyme known to catalyze BHT bioactivation. Further, CYP2B10-catalyzed BHT bioactivation may provide a strategy for future targeted cell ablation technologies or environmental remediation of BHT.

## INTRODUCTION

As a versatile synthetic phenolic antioxidant (SPA), butylated hydroxytoluene (BHT) is widely used in a variety of industrial and commercial products as a preservative including foods, beverages and cosmetics. It is also used to stabilize plastics, rubbers, fuels, oils, and lubricants, while also being added to animal feeds, herbicides and pesticides to prevent their rancidification. Annual global production of BHT is estimated at a staggering 62,000 tons (1). Not surprisingly, BHT and derivatives are readily detectable in marine organisms (2, 3), marine plastics (4), human fingernails (5), urine (6), serum (7) and almost all home dust samples (1, 8). While the FDA considers BHT to be generally recognized as safe (“GRAS”), SPAs such as BHT have recently come under increased regulatory scrutiny in the United States as part of a broader effort to review chemical additives that reach the food supply (9–15).

First described by Marino and Mitchell in 1972 (16), murine intraperitoneal BHT administration results in remarkably cell-specific toxicity to the thin and flat alveolar type 1 (AT1) cells that line the gas-exchanging alveoli. Within 24h of BHT administration, AT1 cells show vacuolization followed by cytoplasmic blebbing and plasma membrane rupture a day later (17). By days 3-4, cuboidal alveolar type 2 (AT2) cells exhibit elongation and differentiation to replace lost AT1 cells (18, 19). In recent years, the BHT model has undergone a maturation from an observed toxicological phenomenon to a mainstay tool for cell biology studies focused on lung epithelial cell repair (20–23). However, the BHT model’s mechanistic basis has remained largely undefined. This knowledge gap surrounding the BHT model has bottlenecked progress with respect to its use in murine injury models, optimizations and translations towards research on human interstitial lung diseases (ILDs).

Enzymatic oxidation of BHT to an electrophilic quinone methide (QM) has been implicated as a mechanism by which BHT can chemically modify protein thiols and amino groups that cause cellular toxicity (24, 25). Pharmacological studies have broadly implicated some type of cytochrome P450 (CYP450) as a driver of BHT enzymatic oxidation and thus BHT toxicity (26), yet no individual CYP450 enzyme has been reproduced to show such BHT activating activity. We viewed this lung-specific knowledge gap as a possible opportunity to discover an enzyme activity responsible for BHT bioactivation into a QM with potentially broad biochemical and health implications.

## MATERIALS AND METHODS

### Materials and reagents

Sources of reagents, antibodies and kits are listed in the supplementary information.

### Analysis of published scRNAseq datasets

Publicly available mouse lung single-cell RNA-seq datasets were analyzed in R using Seurat (v4). For the Strunz et al. whole lung dataset (GEO: GSE141259), raw count matrices (MatrixMarket format) and accompanying gene, barcode, and cell metadata files were downloaded from GEO, imported into R, and used to construct a Seurat object (CreateSeuratObject, min.cells = 3, min.features = 200). Cell identities were assigned using the author-provided cell.type metadata. Mitochondrial content was calculated as the fraction of UMIs mapping to genes prefixed “mt-” (PercentageFeatureSet), and cells were filtered to retain >200 detected genes and <5% mitochondrial reads. Data were log-normalized and scaled with regression of total UMI counts. SCTransform (regressing mitochondrial content) was evaluated but not used for primary analyses due to reduced dynamic range of log2 fold-change estimates.

Curated mouse monooxygenase (n=88) and oxidoreductase gene lists (Swiss-Prot-derived) were imported from CSV files and intersected with dataset features prior to analysis. Differential expression for these gene sets was performed using FindMarkers to test enrichment within specific annotated populations (e.g., AT1 cells, vascular endothelial subsets) compared to all other cells, using min.pct = 0.02 and logfc.threshold = 0 unless otherwise specified. Results were ranked by average log2 fold-change (avg_log2FC) and visualized with custom dot plots encoding effect size, fraction of expressing cells, and adjusted p-values. Cell-type specificity was quantified using a Tau metric computed from mean normalized expression across cell types and visualized alongside enrichment metrics.

### Lentiviral production

Ten cm dishes of HEK293 cells at 80% density were transfected with DNA in a 3:2:1 molar plasmid ratio (transgene : packaging : pseudotype/VSV-G). Per dish, a total of 20 µg DNA with 60 microliters of polyethyleneimine 300k (PEI, 1 mg/mL stock solution) was used. Eighteen hours after transfection, media was changed. Media exchange / collections were performed 1 and 2 days later. Conditioned media was stored at +4 C, pooled, centrifuged at 1000g x 10 min, then pushed through a 0.45 µm PES syringe filter. Conditioned media was stored at +4 C for up to 72 hours then applied to target cells. For long term storage, conditioned media was concentrated using a homemade 3x concentrator (32% PEG-6000, 1.6 M NaCl in PBS) by incubating at +4C overnight, centrifugation at 2000g x 45 min, resuspension in PBS (one hundredth volume of conditioned media) followed by -80C storage.

### Stable transduction of cells

To a single 12 well dish of cells at ∼50% density in 800 µL of media was added 80 µL of lentiviral concentrate and 8 µg/mL polybrene. Three days post transduction, selection was performed with hygromycin 250 µg/mL for at least 7 days (when antibiotic selection was indicated). Cells were kept in 200 µg/mL hygromycin during maintenance culture. Hygromycin was removed (media changed) prior to experiments.

### MTT reduction assay

Thiazolyl Blue Tetrazolium Bromide (MTT) was prepared as 5 mg/mL stock solution in PBS, filter sterilized and stored at -40C until use. Cell media containing FBS was aspirated and a working solution of 0.5 mg/mL MTT in DMEM (without serum) was added to cells for 3 h at 37C followed by 1.8 volumes of isopropanol to help solubilize any precipitated formazan. Absorbance was read at 570 nm.

### Flow cytometry for bystander effect measurements

293T cells stably expressing CYP2B10(WT)-T2A-EGFP or CYP2B10(C436A)-T2A-mCherry were mixed 1:1 and allowed to adhere to tissue culture dishes. When subconfluent, these mixed cell populations were treated in quadruplicate with BHT (50 or 100 µM) or vehicle control for 14 hours before processing for flow analysis. Trypsinized cells were filtered through a 30 µm filter (CellTrics, Sysmex/Partec) and centrifuged before resuspension in PBS containing 2% BSA and stored on ice until analysis. Viability staining was performed using DAPI exclusion, with DAPI added at a final concentration of roughly 100 ng/mL at 15 minutes before analysis. A Sony SH800 flow cytometer was used in analysis mode to analyze the samples and controls. Single-color controls (unstained, GFP-positive only, mCherry-positive only, and a staurosporine-treated DAPI control) were utilized for Sony’s automated compensation routine, with the resultant matrix applied to all test samples. At least 10,000 events from each of the quadruplicate samples were recorded for each sample. Using FlowJo software (version 10.10.1), all compensated samples were analyzed for GFP and mCherry signal after the following gates were applied (in order): scatter gate (FSC-A vs SSC-A), doublet-exclusion gate (SSC-H vs SSC-W), viable cell gate (DAPI-negative) (gating scheme in supplementary information).

### Animal handling and harvesting

All mouse handling was performed in accordance with institutional and IACUC guidelines. Following euthanasia with either cervical dislocation or CO2, mice were perfused through the right ventricle with PBS and lungs harvested. For western blots, lungs were chopped, frozen and stored at -80 C until use. For microscopy, lungs were inflated and fixed in 4% PFA for 2h, then incubated in 30% sucrose overnight followed by cryopreservation in OCT blocks for sectioning.

### Confocal microscopy and quantitative analysis

Slides were subjected to antigen retrieval in citrate buffer pH 6.0 at 95C for 20 min. Tissues were blocked with 2% BSA in PBST for 1h. Details regarding primary antibodies are in the supplementary information. Images were acquired on an Olympus FV4000 confocal microscope using a 40x objective in galvanometer scan mode. Single optical sections (4096 × 4096 pixels at 16-bit) were collected at 4.0 µs/pixel with 2× line averaging in sequential line-scan mode. Four channels were imaged: DAPI (ex 405 nm, em 413–485 nm), Alexa Fluor 488 (ex 488 nm, em 500–540 nm), Alexa Fluor 594 (ex 561 nm, em 583–647 nm), and Alexa Fluor 647 (ex 640 nm, em 650–710 nm). Data were exported in 24-bit RGB and 16 bit raw formats for visual rendering and quantitative analyses, respectively.

For perinuclear analysis of CYP2B10 immunostaining, nuclei were segmented from the DAPI channel and used as masks to measure the mean HOPX immunofluorescence within each nucleus. After subtracting a global background (the median HOPX intensity over all non-nuclear pixels), nuclei were classified as Hopx⁺ (AT1) when the background-subtracted mean exceeded a per-field Otsu threshold scaled by 0.85 (confirmed by visual inspection). All remaining nuclei were pooled as HOPX⁻. A signed Euclidean distance transform was computed from the segmented nuclear boundaries. CYP2B10 was quantified within an asymmetric perinuclear shell extending 2 µm inside and 1 µm outside each segmented nuclear boundary. Intensities were binned (0.25 µm) by signed distance from the boundary and averaged separately based on HOPX status to create a radial profile. Profiles were normalized to that field’s HOPX⁻ far-cytoplasm baseline and averaged across the five fields. Corresponding DAPI profiles were normalized to the interior plateau region.

## RESULTS

### Identification of candidate BHT activating enzymes

Considering the specificity of BHT to cause toxicity to AT1 cells, we hypothesized that BHT toxicity was due to an AT1-cell restricted monooxygenase. Published single cell RNA sequencing (scRNAseq) datasets were utilized to examine the cell-specificity of 88 candidate murine monooxygenase genes. As shown in Figures 1A and 1B, *Cyp2b10* and *Cyp4b1* exhibited AT1 cell enrichment in AT1 cells (27, 28), suggesting them as plausible candidate monooxygenase genes. By examining AT1 cell enrichment alongside Tau score for cell specificity (29), these enzymes demonstrated AT1-specific expression on par with established AT1 cell markers Homeodomain-only protein (*Hopx*), Podoplanin (*Pdpn*) and Receptor for Advanced Glycosylation End-Products (*Ager*) (Figures 1C, 1D).

**Figure 1.**
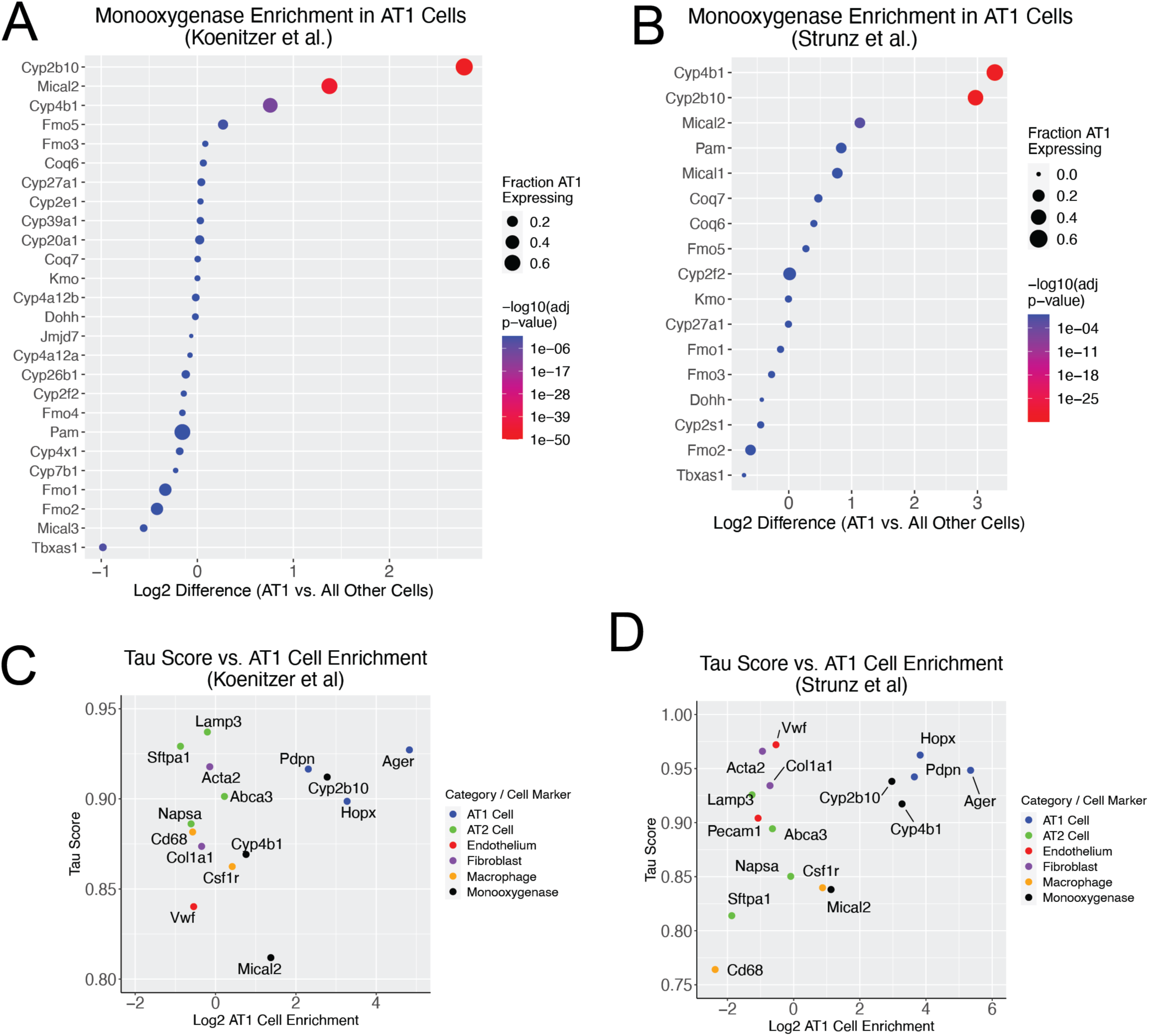
Identification of candidate BHT-activating enzymes. (**A**) Differential expression analysis of monooxygenases in Koenitzer et al. comparing AT1 to all other cells. Of 88 monooxygenases, 70 genes were present in the dataset with 26 genes passing a threshold of 2% of cells expressing each monooxygenase. (**B**) Differential expression analysis of monooxygenases in Strunz et al. comparing AT1 to all other cells. Of 88 monooxygenases, 45 genes were present in the dataset with 17 genes passing a threshold of 2% of cells expressing each monooxygenase. (**C**) Dotplot of relative expression from Strunz et al. for established AT1 cell marker *Hopx*, candidate monooxygenases *Cyp2b10*, *Mical2* and *Cyp4b1*, and ubiquitously expressed beta-actin. Scatterplot of Tau score vs. AT1 cell enrichment from (**D**) Koenitzer et al. and (**E**) Strunz et al. The three top candidate monooxygenases (*Cyp2b10*, *Cyp4b1*, *Mical2*) are in black with each color representing established cellular markers. Higher Tau score indicates increasing restriction of gene expression and thus increasing overall cellular specificity. As expected, AT1 cell markers (blue) exhibit high Tau score and AT1 cell enrichments. The AT1 cell markers are accompanied by *Cyp2b10* in both datasets.

Both *Cyp4b1* and *Cyp2b10* are established transcriptional markers of terminal AT1 cell differentiation (30), yet CYP4B1 inhibitors fail to protect against BHT-induced lung injury (26). Thus we focused on whether CYP2B10 protein is expressed in AT1 cells. Analysis of healthy mouse lung tissue by immunofluorescence using a validated anti-CYP2B antibody (Figure S1) revealed AT1-specific expression of CYP2B10 in perinuclear distribution consistent with expected localization to the cytoplasmic side of the endoplasmic reticulum (Figure 2A). Radial profiling of nuclei based on HOPX status revealed increased CYP2B10 intensity in the immediate perinuclear zone restricted to HOPX+ nuclei (Figure 2B). Analysis of CYP2B10 expression by western blot showed an increase from postnatal day 6 to 28 (Figure 2C) consistent with the established timeframe when mice acquire BHT sensitivity (31). These data also establish CYP2B10 as a novel protein marker for the AT1 cell body (compared to traditional pan-membrane markers), which to our knowledge has not been previously described.

**Figure 2.**
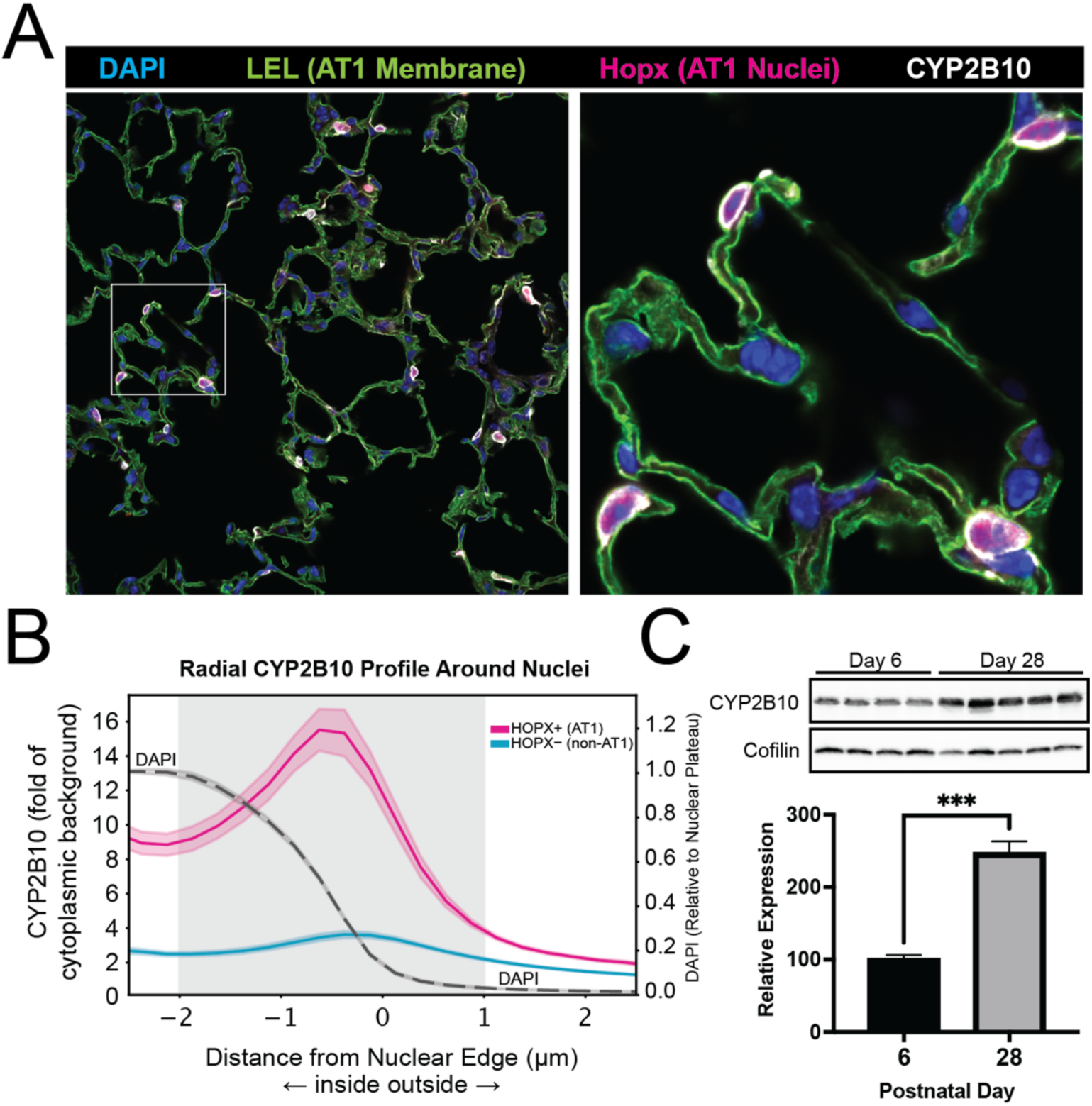
Cytochrome P450 2B10 is expressed in murine alveolar type 1 cells and increases during postnatal period. (**A**) Healthy mouse lung from strain C57BL/6 was harvested, cryosectioned and stained for CYP2B10, AT1 nuclear marker HOPX, AT1 membrane-reactive Lycopersicon Esculentum Lectin (LEL) and DAPI. Confocal microscopy was performed at 40x magnification. Data are representative of n = 3 mice. (B) Radial profile of CYP2B10 intensity based on HOPX status. Mean CYP2B10 intensity plotted as a function of distance from the segmented nuclear boundary (x = 0). The grey band marks the 2 µm-in/1 µm-out perinuclear quantification zone. The dashed grey curve shows the DAPI profile. Data are mean ± SEM across five 40x fields. Total nuclei quantitated are 97 and 1047 for HOPX+ and HOPX-, respectively. (**C**) Western blot of CYP2B10 expression based on postnatal age. Mouse lungs were harvested at the indicated postnatal age (n = 4 for day 6, n = 5 for day 28) with western blotting for CYP2B10 and cofilin as loading control. Using densitometry, CYP2B10 expression increased approximately 2.5-fold from day 6 to day 28 (unpaired t-test with Welch’s correction, t = 8.330, df = 5.062, \*\*\**p* = 0.0004). Data are mean ± SEM.

### Assessment of BHT sensitivity in cells stably expressing Cyp4b1 or Cyp2b10

Murine *Cyp2b10* and *Cyp4b1* were amplified from whole mouse lung cDNA (Figure S1). The cloned *Cyp2b10* cDNA yielded a 491 amino acid ORF identical to UniProtKB-TrEMBL entry Q9WUD0. The cloned ORF for *Cyp4b1* was identical to UniProtKB entry Q64462 bearing 511 amino acids. These ORFs were subcloned into a lentiviral expression vector and used to generate stable *Cyp4b1*- and *Cyp2b10*-expressing cells. As shown in Figure 3A, parental HEK293T cells and those expressing CYP4B1 exhibited no change in cellular viability when challenged with BHT. However, expression of CYP2B10 resulted in pronounced BHT sensitivity, while expression of the enzymatically inactive C436A mutant of CYP2B10 (lacking the heme axial thiolate) showed no sensitivity to BHT. In an analogous approach, the effect of CYP450 inhibitors was explored in cultured NIH-3T3 fibroblasts stably expressing wild-type or C436A mutant of CYP2B10. Cells were pre-treated for 1 hour with SKF-525a or 1-aminobenzotriazole (ABT), exposed to vehicle (EtOH) or 50 µM BHT for 18 h followed by viability assessment with cell counting. Both CYP450 inhibitors protected cells from BHT-induced cell death (Figure 3B) with ABT able to provide complete protection. Collectively, these results directly implicate CYP2B10 enzymatic activity in BHT-induced cell death and establish this mechanism is operative in human and mouse cell lines.

**Figure 3.**
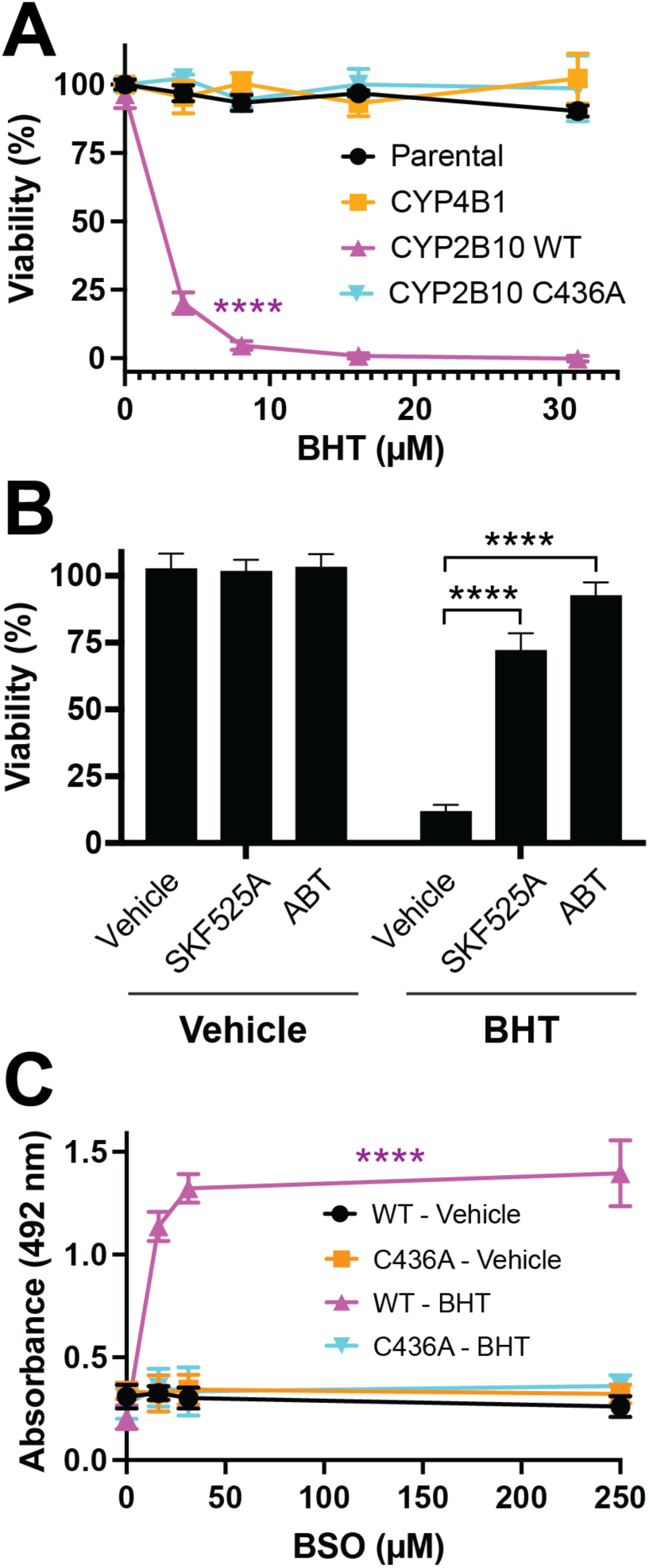
Cultured cells expressing enzymatically active CYP2B10 are sensitized to BHT and protected by endogenous GSH. (**A**) Parental HEK293T cells or HEK293T cells stably expressing CYP4B1, CYP2B10 WT or CYP2B10 C436A inactive mutant were treated with varying concentrations of BHT for 18h followed by viability assessment via MTT reduction assay. Points show mean ± SD (n = 3 for all conditions except CYP2B10 WT n = 5). Fitting a nonlinear four variable curve to CYP2B10 data points shows IC50 = 2.3 µM (95% CI 1.66 to 2.86). (**B**) HEK293T cells stably expressing CYP2B10 WT were pre-treated for 1 h with vehicle (DMSO), 20 μM SKF-525a or 1 mM 1-aminobenzotriazole (ABT), then exposed to vehicle (EtOH) or 50 μM BHT for 18 hours. Viability was assessed using trypan blue exclusion. Data are mean ± SD, n = 4. Both inhibitors protected against BHT (two-way ANOVA, treatment × inhibitor interaction F(2,18) = 135.2, *p* < 0.0001; Dunnett’s post hoc test, \*\*\*\**p* < 0.0001). (**C**) HEK293T cells expressing WT or C436A CYP2B10 were treated with varying concentrations of L-Buthionine-(S,R)-Sulfoximine (BSO) for 24h followed by vehicle or 5 µM BHT for 4h. Media was collected for LDH release assay. Data are mean ± SD, n = 3. Two-way ANOVA revealed a genotype/treatment × BSO interaction (F(9,32) = 36.98, *p* < 0.0001). Dunnett’s post hoc test for each group vs 0 µM BSO (**** *p* <0.0001). All comparisons in vehicle and C436A-BHT groups were non-significant.

### Glutathione protects from CYP2B10-catalyzed BHT toxicity

As electrophilic and unstable species, QMs are expected to react with a range of cellular nucleophiles including glutathione (GSH). To assess the presence of an electrophilic species as the mediator of BHT-induced cell death, we hypothesized that lowering of cellular glutathione would further sensitize cells to BHT in a CYP2B10-dependent manner. As anticipated, HEK293T cells expressing CYP2B10 were unaffected by overnight exposure to γ-glutamylcysteine synthetase inhibitor buthionine sulfoximine (BSO). However, when these same cells were treated with BSO, low micromolar concentrations of BHT lead to rapid detachment, shrinkage and release of LDH into the culture medium (Figure 3C). In contrast, cells expressing inactive CYP2B10 (C436A) did not exhibit synergy between BHT and BSO. These findings strongly suggest that CYP2B10 mediates the production of an electrophilic BHT derivative (i.e., some type of QM) and that GSH is the cell’s primary protection mechanism against downstream BHT electrophilic injury.

### BHT-Driven Cell Death is Cell-Autonomous and Lacks Bystander Effect

In mice, BHT exhibits highly selective cytotoxicity towards AT1 cells suggesting cell-restricted mechanisms for chemical activation and toxicity. To probe this specific hypothesis, HEK293T cells expressing CYP2B10(WT)-T2A-EGFP and CYP2B10(C436A)-T2A-mCherry were co-cultured 1:1, exposed to vehicle or 50 µM BHT for 14 h, then assessed for toxicity to bystander cells (i.e., mCherry+ population). Exposure to BHT resulted in rounding, detachment and global death of WT enzyme-expressing cells while adjacent C436A-expressing cells showed no morphological changes suggestive of injury (Figure 4A). Flow cell cytometry (Figure 4B) showed BHT significantly increased mCherry-positive cells (mean 47.3% → 83.8%) with a concomitant decrease in the EGFP-positive population (mean 44.0% → 1.7%). The persistence of C436A-expressing cells adjacent to dying WT cells suggests that BHT toxicity is cell-autonomous without detectable bystander effect. These findings are consistent with the phenotypic properties of the BHT murine model and suggest that ectopic CYP2B10 expression + BHT might be a useful tool for ablating specific cellular subpopulations while sparing neighboring cells.

**Figure 4.**
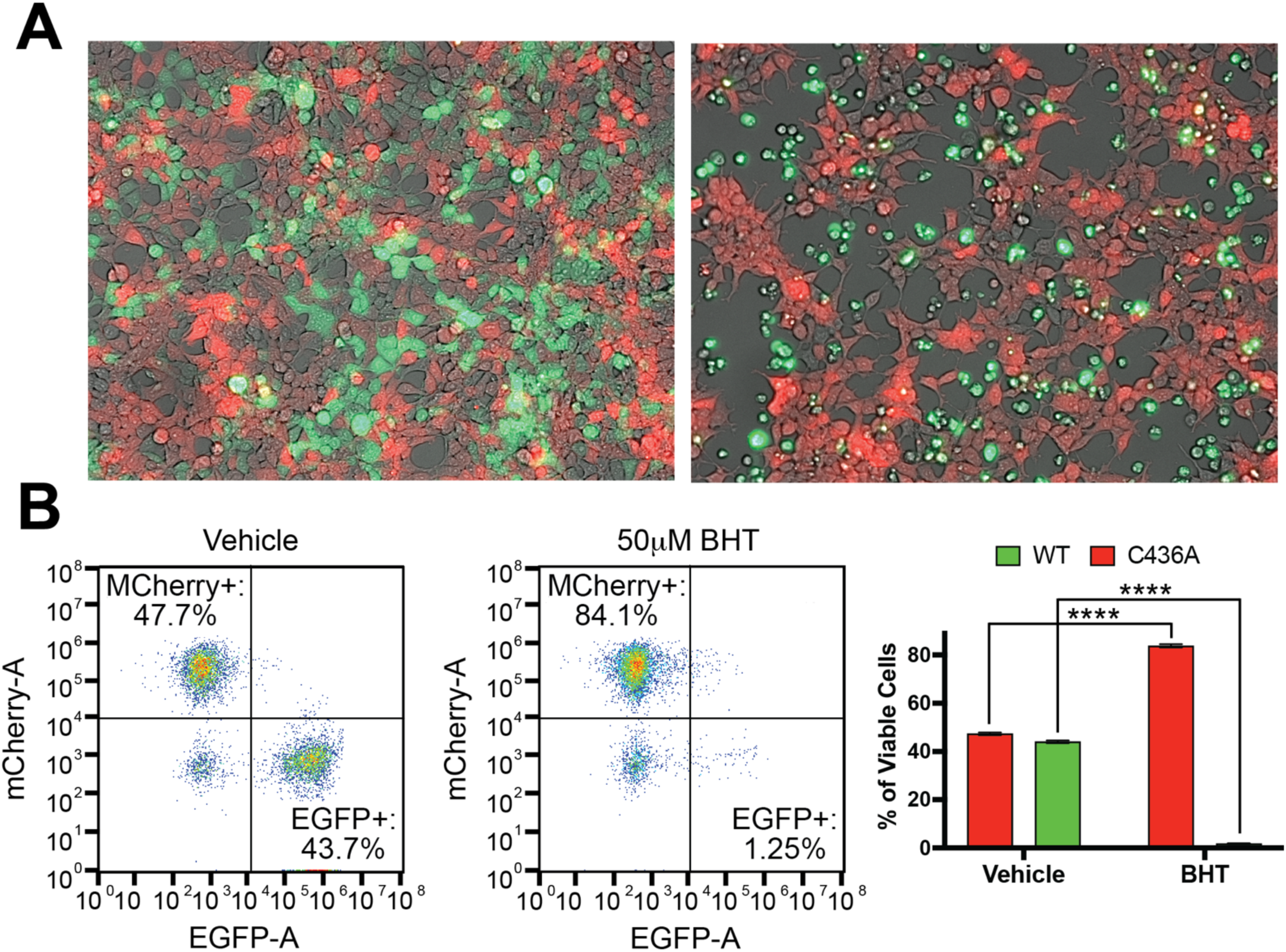
BHT induces death in a cell-autonomous manner. HEK293T cells stably expressing wild-type CYP2B10-T2A-EGFP or C436A CYP2B10-T2A-mCherry were co-cultured 1:1 and exposed to vehicle or 50 µM BHT for 14 h. Cells were analyzed by (**A**) fluorescence and brightfield microscopy at 10x magnification and (**B**) fluorescence-based flow cytometry. Two-way ANOVA: genotype × treatment interaction F(1,12) = 22609, *p* < 0.0001 with Šídák’s multiple comparison correction, **** *p* < 0.0001. Data are mean ± SD, n = 4.

### BHT Bioactivation Triggers Hallmarks of Apoptosis and DNA Damage Response

To understand the molecular events underpinning CYP2B10-catalyzed cell death, HEK293T cells were subjected to a panel of western blots in response to BHT exposure. As shown in Figure 5A, cells expressing functional CYP2B10 exhibited time-dependent activation of stress kinases including phosphorylation of p38 and c-Jun N-terminal kinases at Thr^180^/Tyr^182^ and Thr^183^/Tyr^185^, respectively. Transcription factor c-Jun was phosphorylated on Ser^73^ and exhibited a moderate degree of stabilization (increased expression), which together are suggestive of AP-1 activation in response to BHT activation.

**Figure 5.**
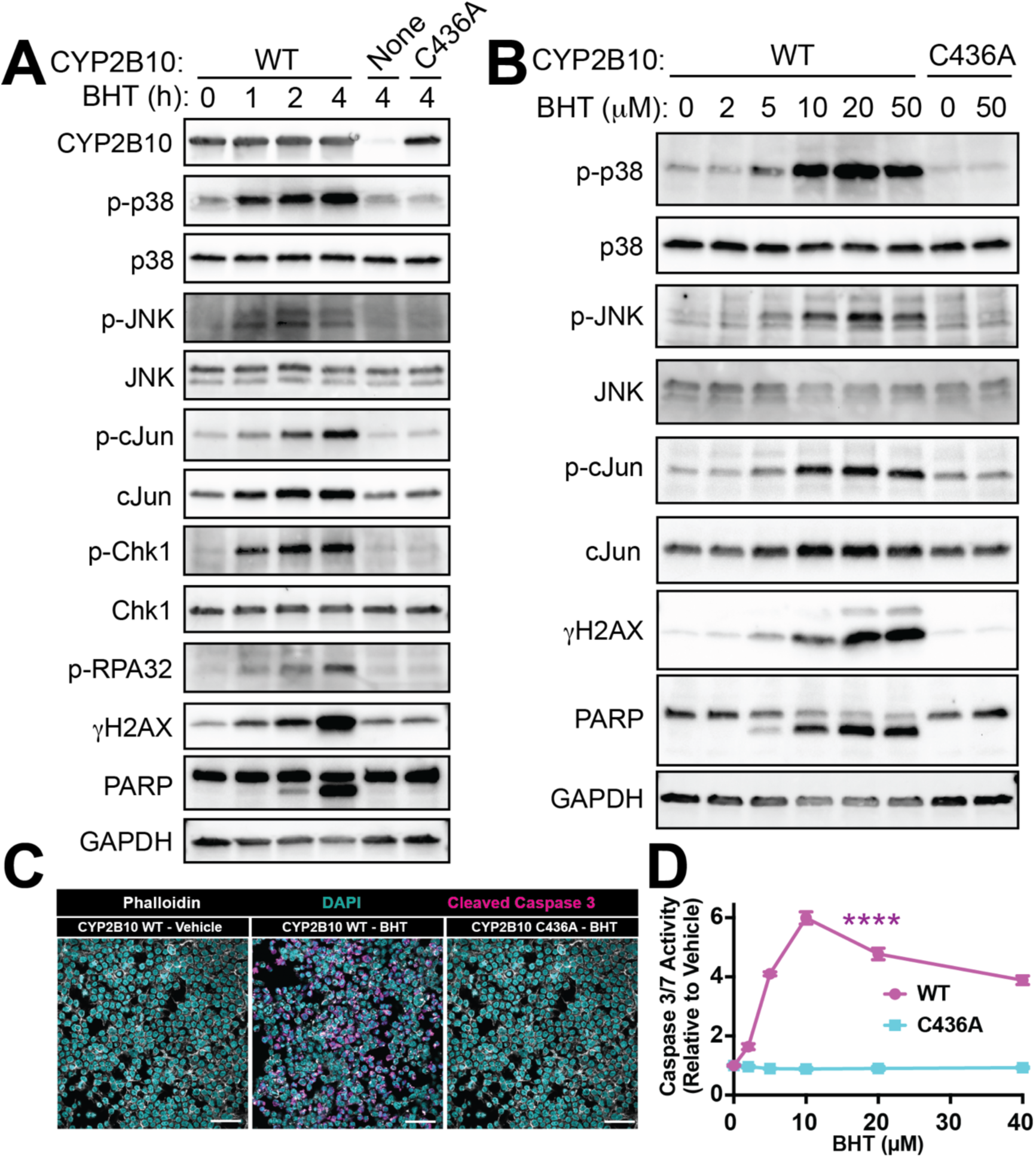
Bioactivation of BHT leads to stress kinase activation, DNA damage response and intrinsic apoptosis in an enzyme-dependent manner. (**A**) Time course of cellular events after BHT exposure. Parental HEK293T cells or those expressing CYP2B10 WT vs inactive C436A mutant were exposed to vehicle (EtOH) or 20 µM BHT for indicated duration, collected, lysed and subjected to western blot for the indicated proteins. (**B**) NIH-3T3 fibroblasts expressing CYP2B10 WT or inactive C436A mutant were exposed to vehicle or BHT at indicated concentrations for 12h, collected, lysed and subjected to western blot for indicated proteins. (**C**) HEK293T cells expressing WT or C436A CYP2B10 were exposed to vehicle or 20 uM BHT for 2.5h, fixed with PFA, immunostained for active/cleaved Caspase-3 and imaged by confocal microscopy at 40x magnification. Scale bar = 50 µm. (A), (B) and (C) are representative of at least n = 3 separate experiments. (**D**) HEK293T cells expressing WT or C436A CYP2B10 were treated with vehicle or indicated concentrations of BHT for 3.5h and subjected to Caspase-3/7 Glo luminescence assay. Two-way ANOVA: genotype × BHT interaction F(5,60) = 1052, *P* < 0.0001; Dunnett’s multiple comparisons vs 0 µM BHT, \*\*\*\**P* < 0.0001 (WT at all concentrations). All C436A comparisons were non-significant. Data are mean ± SD, n = 6.

With respect to the possibility of DNA damage, a quinone methide derivative of BHT would be expected to alkylate DNA without directly promoting crosslinking or double strand breaks. Therefore, we hypothesized that ataxia telangiectasia and Rad3-related (ATR) kinase may be activated by BHT, leading to phosphorylation of checkpoint kinase 1 (Chk1). As shown in Figure 5A, Chk1 is phosphorylated on Ser^345^ in a similar timeframe as stress kinases (p38, JNK), suggesting rapid cell cycle arrest in a CYP2B10-dependent manner. Phosphorylation of replication protein A (RPA32) on Ser^33^ and histone H2A.X on Ser^139^ (γH2AX) were also observed, indicative of a global DNA damage response triggered only by the combination of active CYP2B10 and BHT. Following the wave of stress kinases and DNA damage response was cleavage of Poly ADP-Ribose Polymerase (PARP), indicative of executioner caspase activity and suggestive of intrinsic apoptosis.

To determine whether this was a phenomenon restricted to human epithelial cell line (i.e., HEK293T), complementary experiments were undertaken in murine NIH-3T3 mesenchymal cell line. As shown in Figure 5B, NIH-3T3 cells expressing enzymatically active CYP2B10 demonstrated a similar response to BHT, whereas those expressing the inactive C436A mutant were unaffected by BHT up to its aqueous solubility limit of ∼50 µM. These cells also showed dose-dependent cleavage of PARP suggestive of caspase-3 activation. To directly probe the role of executioner caspase activity, CYP2B10-expressing HEK293T cells were subjected to immunofluorescence for cleaved caspase 3 (Figure 5C) and chemiluminescent detection of caspase-3 substrate cleavage (Figure 5D). In both cases, caspase-3 is rapidly activated and requires the presence of both BHT and enzymatically active CYP2B10. As shown in Figure 5D, caspase-3 activation showed a biphasic response to BHT, peaking around 10-20 µM. Higher concentrations exhibited less caspase-3 activation despite prominent cell death and LDH release, overall suggesting a situation where massive BHT-induced damage triggers less-regulated cell death mechanisms like necrosis.

### LC-MS/MS Identifies the Addition of t-Butyl Hydroxylated BHT (BHTOH) to Protein Thiols

Given the likely mechanism of electrophilic injury, we hypothesized that CYP2B10 would indirectly promote alkylation of intracellular proteins through activating BHT into a quinone methide. To test this theory, NIH-3T3 fibroblasts expressing either WT or C436A CYP2B10 were exposed to 20 µM BHT for 12 hours, lysed, digested and analyzed by LC-MS/MS (bottom up proteomics). To allow for a broad and unbiased analysis of CYP2B10-catalyzed protein modifications, results from data dependent acquisition (DDA) were subjected to an open search algorithm (32) capable of comprehensively surveying mass shifts from -150 to 500 Da (33). Analysis of 150 matched mass shifts (each with at least 5 PSMs in one condition) demonstrated the presence of a dominant +234.162 Da modification in cells expressing WT CYP2B10 exposed to BHT but not the same cells exposed to vehicle or expressing inactive C436A CYP2B10 (Figures 6A, 6B, Table S1). This mass shift corresponds to Michael addition of t-butyl hydroxylated BHT (BHTOH, Unimod entry 498). Open searches did not reveal Michael addition of BHT at the expected +218.167 Da (Unimod entry 176). These data strongly implicate a two-step bioactivation process involving t-butyl hydroxylation (to BHTOH) followed by oxidation to a quinone methide (BHTOH-QM), which is consistent with *in vivo* and *ex vivo* studies suggesting a correlation between BHT t-butyl hydroxylation and pneumotoxicity in mice (34–36).

**Figure 6.**
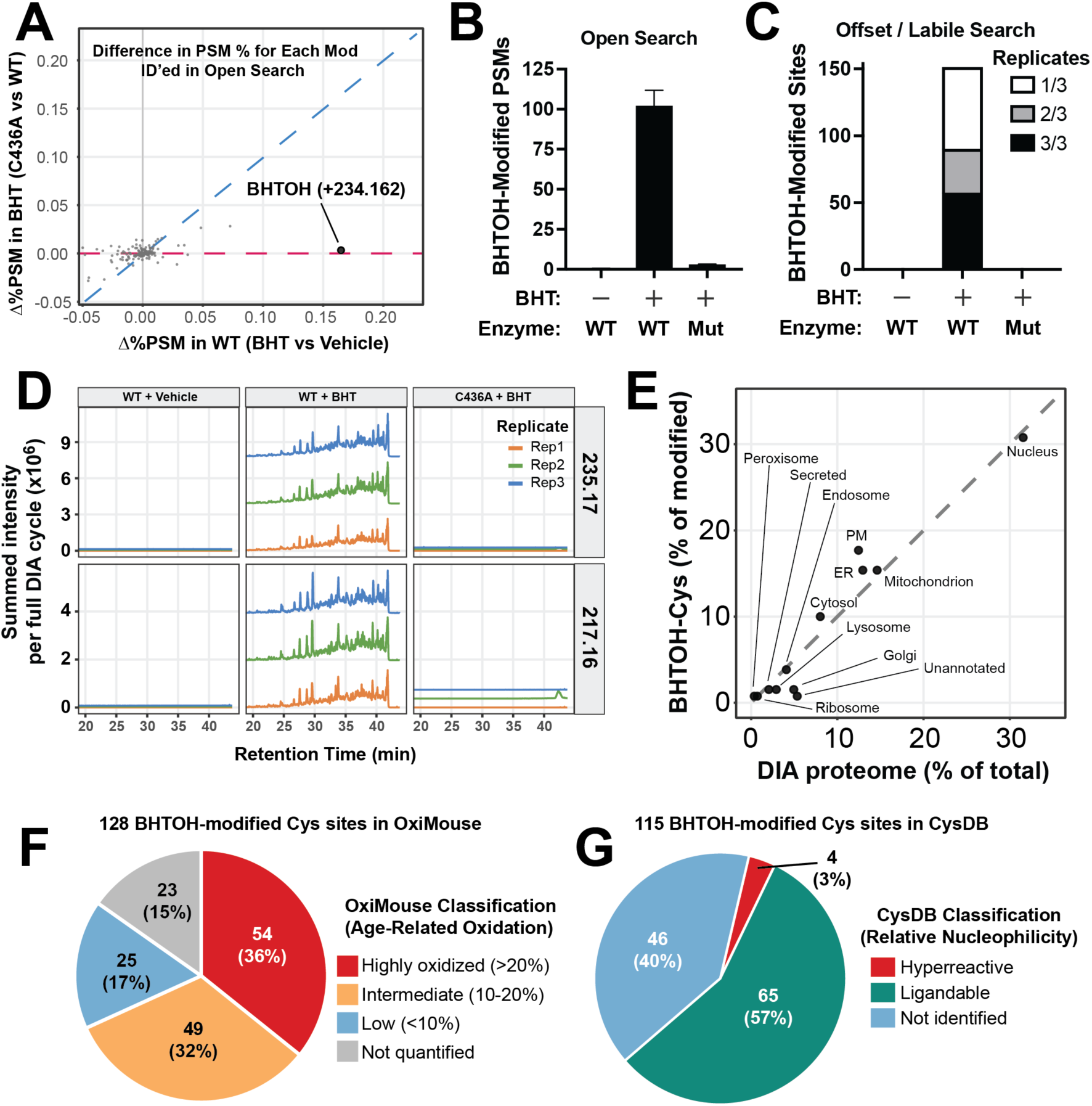
Characterization of protein Cys alkylation during BHT-bioactivation. NIH-3T3 cells stably expressing CYP2B10 WT or C436A were exposed to vehicle or 20 µM BHT for 12h, collected, lysed, trypsinized and subjected to LC-MS/MS. Three biological replicates were analyzed for each condition (WT + vehicle, WT + BHT, C436A + BHT). (**A**) Results of open search shown as scatterplot where each point represents a distinct modification. Shown are the 150 modifications for which at least one condition had ≥ 5 PSMs. Units are difference in percentage of total PSMs for BHT vs. Vehicle (x-axis) and C436A vs. WT (y-axis). Modifications that are “catalysis-dependent” exhibit high x-axis and low y-axis values, respectively. Shown in bold is the BHTOH modification. (**B**) Open search results from DDA data for total number of BHTOH-containing PSMs in each condition. (**C**) Number of BHTOH-modified sites from labile/offset search. For (B) and (C), WT = wild type, Mut = C436A inactive mutant. (**D**) Extracted ion chromatograms (XICs) for BHTOH-derived diagnostic ions at 235.17 and 217.16 m/z from DIA data. (**E**) Assessment of BHTOH modification by subcellular localization. Scatterplot showing percentage of subcellular annotations for BHTOH-modified sites on y-axis and total proteome representation on x-axis. Each point is a distinct subcellular annotation. Distance from dashed line (slope = 1) indicates deviation in BHTOH-modified sites compared to reference proteome. (**F**) Comparison of BHTOH-modified sites to OxiMouse database of redox-active Cys sites. (**G**) Comparison of BHTOH-modified sites to CysDB database of nucleophilic / ligandable Cys sites.

Benzylic adducts of BHT are known to be highly unstable during ionization and fragmentation methods employed in LC-MS/MS (25, 37), likely due to resonance stabilization of benzyl cations formed during heterolytic S-C bond cleavage. This property poses a barrier for mapping sites of BHT-QM (or BHTOH-QM) alkylation by traditional variable search methods that rely on adduct stability during ionization and fragmentation (24). Consistent with the lability of BHT-adducts, manual review of MS/MS spectra from open searches showed prominent diagnostic ions at m/z 235.17 and 217.16 corresponding to loss of BHTOH and BHTOH minus H2O, respectively, from modified peptides. These diagnostic ions were employed in an offset/labile algorithm (38, 39) on DDA data to identify a total of 151 sites of Cys alkylation with BHTOH on 130 distinct proteins (Figure 6C, Table S2). Searching with Lys or His as sites of modification did not reveal additional sites of BHTOH modification despite such amino-adducts showing higher tolerance of fragmentation methods compared to Cys thiol-adducts (25). Examination of data independent acquisition (DIA) data through extracted ion chromatography (XIC) for BHTOH-reporter ions confirmed the lability of this modification (Figure 6D, Table S3). Collectively these data implicate Cys thiols as the dominant site of protein alkylation when BHT is bioactivated to BHTOH-QM. Further, Cys-BHTOH adduct lability likely limits our identifications to only a subset of truly modified proteins.

Given the chemical uniqueness of BHT bioactivation, we sought to understand whether BHTOH-modified sites exhibit unique biochemical or cellular properties. To determine if protein subcellular localization determines reactivity towards BHTOH-QM, the subcellular annotations of BHTOH-modified proteins were compared to the reference proteome (Figure 6E). Plasma membrane associated proteins showed a modest 5.2% enrichment over total proteome abundance. However no subcellular localization achieved statistical significance when corrected for multiple hypothesis testing. Of these 151 BHTOH-modified sites, 128 sites (85%) are listed in OxiMouse (40) as showing some degree of age-dependent oxidation (Figure 6F), suggesting that BHTOH-QM is reacting with known nucleophilic and chemically accessible Cys thiols. A well-studied example is Cys113 on p62 (Sqstm1), which is a nucleophilic / redox-active site that allows cellular activation of autophagy in response to oxidative stress (41).

The CysDB is a comprehensive database of 64,681 human Cys sites obtained from 9 published proteomic datasets, of which 15.8% are considered “ligandable” and 0.8% are considered “hyper-reactive” indicating a high propensity for quantitative alkylation (42). To facilitate matching of BHTOH-modified sites to CysDB, the 151 BHTOH-modified sites were mapped to 118 orthologous human Cys sites. Of these, 115 (97%) were listed in CysDB, including 69 (60%) that are ligandable and 4 (3.5%) that are hyper-reactive sites (Figure 6G). These correspond to relative enrichment factors of 3.8x and 4.4x for ligandable and hyperreactive sites, respectively. These results again show a propensity of BHTOH-QM to react with known nucleophilic Cys thiols. However, 46 (40%) of the BHTOH-modified Cys sites are not considered ligandable in CysDB, suggesting that BHTOH-QM also alkylates Cys sites not accessible to typical exogenous alkylating agents that comprise the bulk of CysDB (i.e., iodoacetamides, chloroacetamides, acrylamides). This discrepancy may be attributed to the *in situ* conversion of BHT into BHTOH-QM by CYP2B10 within living cells compared to the extracellular origin of alkylating agents employed in CysDB.

### CYP2B10-Catalyzed BHT Activation Triggers Global Proteome Changes Consistent with Electrophilic Injury, Cell Cycle Arrest and Programmed Cell Death

To maximize proteome depth and support functional analyses, lysates from BHT-treated NIH-3T3 cells were subjected to data independent acquisition (DIA) LC-MS/MS. Across three biological replicates, 8304 total proteins were quantitated (Table S4). As shown in Figure 7A, activation of stress-related signaling pathways were readily apparent. These include BHT-upregulation of Nrf2 targets such as glutamate-cysteine ligase (Gclc), cystine-glutamate transporter (Slc7a11), glutathione S-transferase A2 (Gsta2), ferritin light chain 1 (Ftl1), ferritin heavy chain (Fth1) and heme oxygenase 1 (Hmox1). Consistent with this trend, Keap1 – the master negative regulator of Nrf2 – is BHT-downregulated (Log2FC - 1.37). Likewise, proteins involved in DNA damage and unfolded protein responses (UPR) showed upregulation, while those involved in cell cycle progression were downregulated. Across the 8304 quantified proteins, 1821 showed statistically significant change in response to BHT (|log₂FC| > 0.5, adj *p* value < 0.05). As shown in Figure 7B, 494 proteins (5.9% of the proteome) and 1327 (16.0% of the proteome) were up- and downregulated, respectively. These BHT-induced proteome changes were overwhelmingly dependent on catalytically active CYP2B10 (Figure 7C) with only 1% of the BHT-responsive proteome showing preservation in enzymatically inactive (CYP2B10 C436A) cells. As shown in Figure 7D, a BHT-responsive trend for enzymatically active CYP2B10 is observed across the entire proteome.

**Figure 7.**
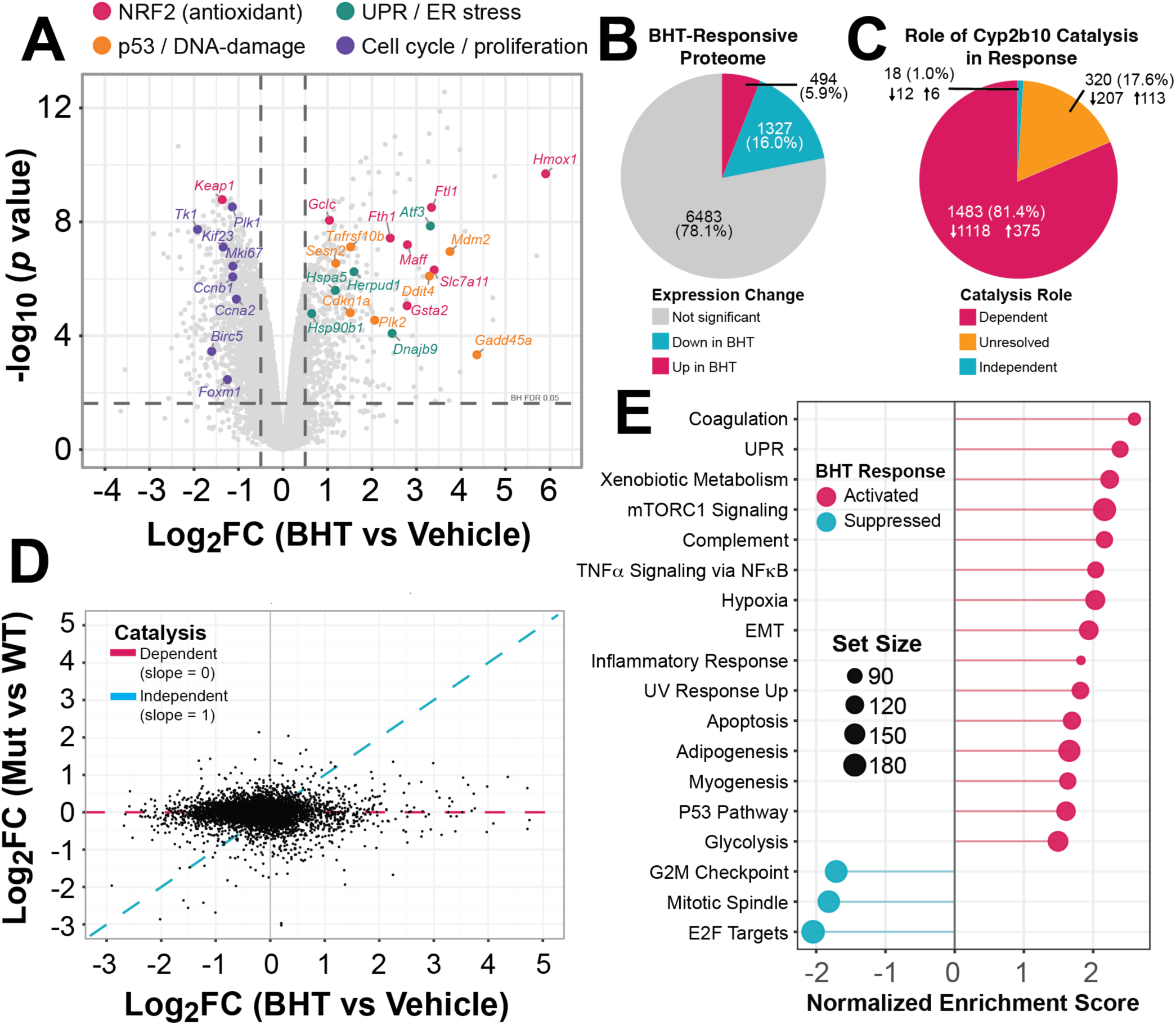
Assessment of proteome-wide expression changes during BHT-induced cell death. NIH-3T3 cells stably expressing CYP2B10 WT or C436A were exposed to vehicle or 20 µM BHT for 12h, collected, lysed, trypsinized and subjected to LC-MS/MS (n = 3 for each condition). (**A**) Volcano plot of BHT-responsive proteome within CYP2B10 WT-expressing cells. Highlighted are four functional categories with up to 8 representative proteins reaching statistical significance (|log₂FC| > 0.5, adjusted *p*-value < 0.05). (**B**) Pie chart of BHT responsive proteome. Proteins were considered responsive to BHT if they changed significantly in WT condition (-/+ BHT) with |log₂FC| > 0.5 and BH-adjusted *p* < 0.05. Non-significant (n = 6,483). (**C**) Catalysis dependence for BHT-responsive proteome. The 1,821 responders were subcategorized using the catalytic-dead C436A mutant in three pairwise limma contrasts (empirical-Bayes moderated t, BH-FDR within contrast). Catalysis categories are the intersection of these per-contrast calls; non-significance is used as a (power-limited) proxy for equivalence, so intermediate and underpowered patterns are pooled as “unresolved”. (**D**) Scatterplot of catalysis across entire proteome. Each point is one protein (n = 8,304). The x-axis represents the BHT response with catalysis intact (WT cells, BHT vs Vehicle) while y axis represents catalysis abolished (C436A+BHT vs WT+Vehicle). Proteins on the magenta line (y = 0) lose their response in the catalytic-dead mutant and are catalysis-dependent, whereas proteins on the cyan line (y = x) respond equally with or without catalysis (i.e., catalysis-independent). A linear fit of y on x gives a slope of 0.038, nearly parallel to the magenta line. The fit line is not shown for clarity. (**E**) Gene Set Enrichment Analysis (GSEA) using clusterProfiler with MSigDB mouse Hallmark sets. Shown are the top 15 activated and all suppressed gene sets ranked by adjusted *p* value. Only 3 gene sets reach FDR < 0.05 for suppressed. GSEA abbreviations: UPR, unfolded protein response; EMT, epithelial mesenchymal transition.

Gene set enrichment analysis (GSEA) using the MSigDB Hallmark gene set (43) showed upregulation of stress-related pathways such as unfolded protein response, xenobiotic metabolism, mammalian target of rapamycin (mTOR), apoptosis and p53 signaling. Within Hallmark, only three gene sets met statistical significance for BHT-induced downregulation. These included G2M checkpoint, mitotic spindle, and E2F targets, which collectively fit with a program of cell cycle arrest and blunting of replication. In summary, the GSEA results (Table S5) are consistent with the targeted proteomic analyses (Figure 7A) and western blots focused on key stress response pathways (Figures 5A/B).

### Structure Activity Relationship Reveals Essential Elements for CYP2B10-Bioactivation of p-Cresol Analogues

To gain an understanding of the chemical determinants necessary for BHT bioactivation, analogues of BHT (Figure 8A) were applied to HEK293T cells expressing CYP2B10 followed by assessment of cellular toxicity by LDH release. As shown in Figure 8B, removal of one ortho t-butyl group from BHT (Compound A) preserved cellular toxicity comparable to BHT. However, isomerization of the para methyl group to the ortho position (Compound B) or replacement of the single remaining ortho t-butyl group with a methyl group (Compound C) led to a dramatic loss of cytotoxicity. Lengthening the 4-alkyl substitution from a methyl to ethyl moiety (Compound D) resulted in a modest decrease in toxicity. Collectively these findings demonstrate the requirement of a single t-butyl group in the ortho position and an alkyl substituent (preferably methyl) in the para position. These chemical requirements are consistent with a chemical bio-activation process of t-butyl hydroxylation followed by aromatic oxidation to a quinone methide (e.g. BHTOH-QM), with slightly diminished chemical reactivity when the methide is substituted with an alkyl group (Compound D).

**Figure 8.**
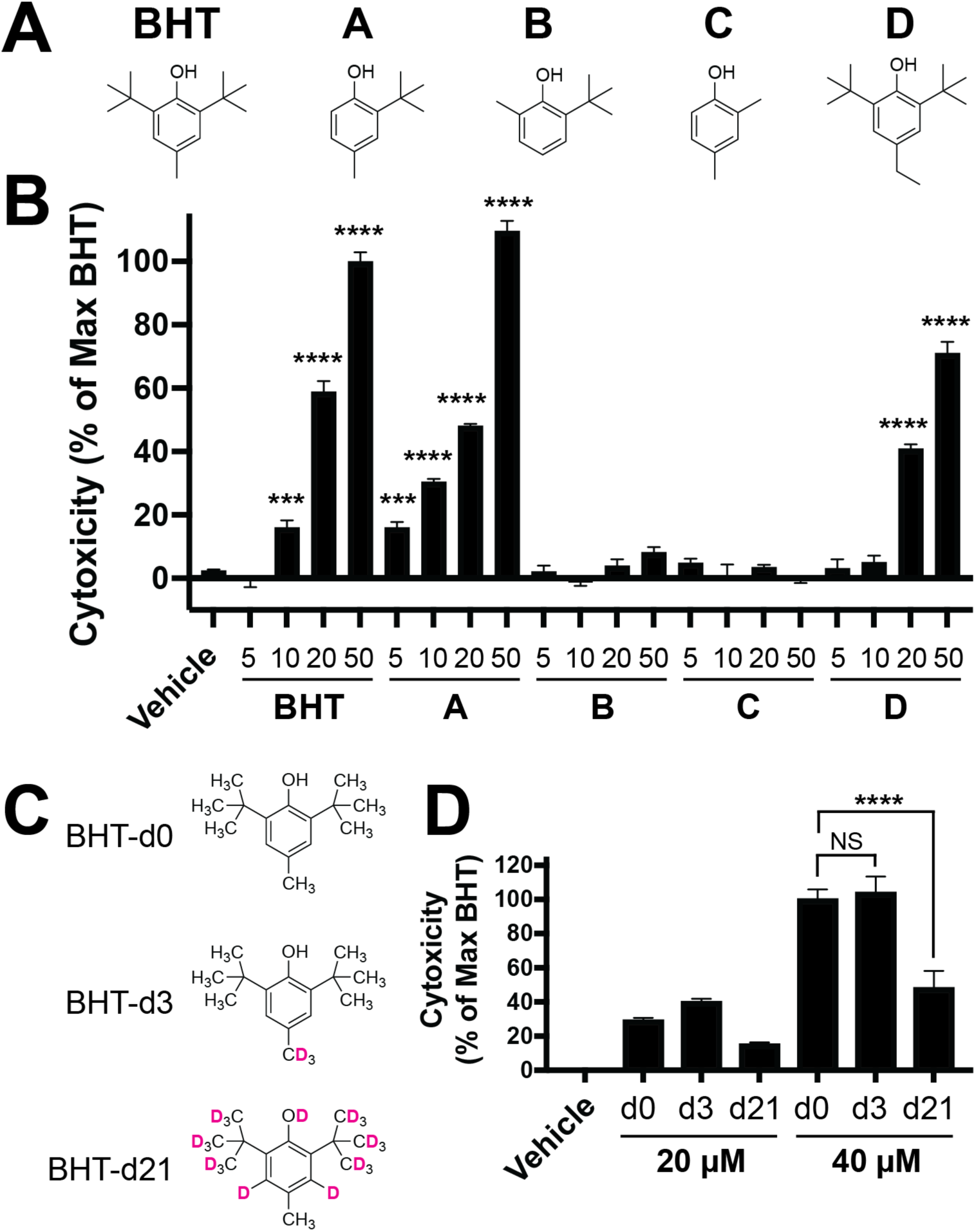
Structure-activity relationship and isotopic effects on enzyme-mediated BHT bioactivation. (**A**) BHT analogues tested: BHT = 2,6-Di-*t*-butyl-4-methylphenol, A = 2-*t*-Butyl-4-methylphenol, B = 2-*t*-Butyl-6-methylphenol, C = 2,4-dimethylphenol, D = 2,6-Di-*t*-butyl-4-ethylphenol. (**B**) Cytotoxicity as measured by LDH release after 8 h incubation with indicated analogue concentration. One-way ANOVA (F(20, 63) = 235.1, P < 0.0001) with Dunnett’s post hoc test of treatments vs vehicle, \*\*\**P* < 0.001, \*\*\*\**P* < 0.0001. Data are mean ± SD, n = 4. (**C**) Isotopomers of BHT include BHT-d3 (deuterium on para-methyl group) and BHT-d21 (all hydrogens replaced with deuterium except for the para-methyl group). (**D**) Cytotoxicity as measured by LDH release after 5 h incubation with vehicle, BHT or indicated isotopomer. Two-way ANOVA: deuterium label × dose interaction F(2,18) = 5.17, *P* = 0.017 with Dunnett’s post hoc test of BHT-d3 and BHT-d21 vs unlabeled (BHT-d0) within each dose, \*\*\*\**P* < 0.0001, NS = not significant. Data are mean ± SD, n = 4.

**Figure 9.**
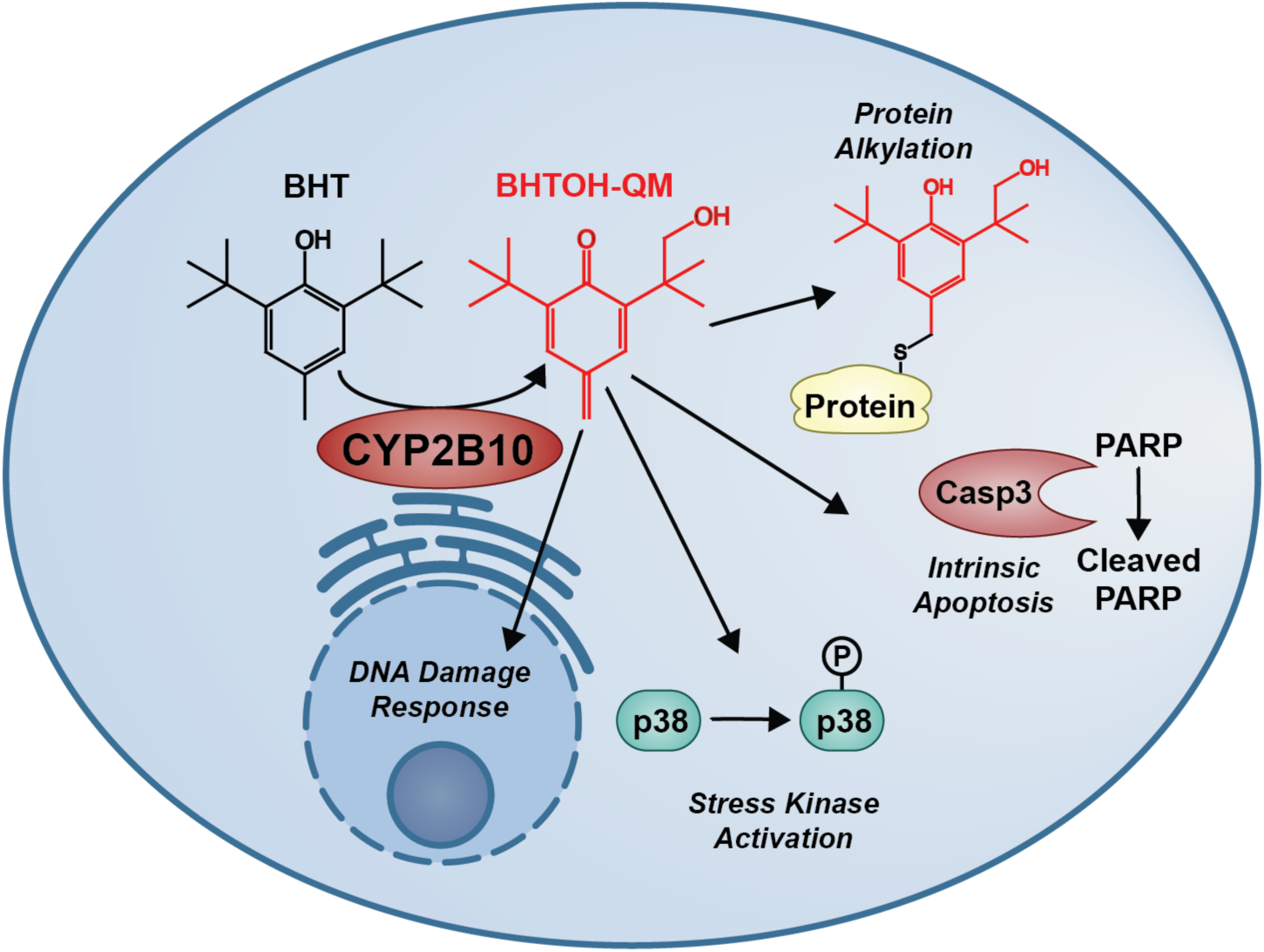
Model of BHT bioactivation and downstream cellular consequences. By virtue of its lipophilicity, BHT freely diffuses into the cell. Upon encountering CYP2B10 at the cytoplasmic surface of the endoplasmic reticulum, BHT undergoes hydroxylation (likely on a single *t*-butyl group) to BHTOH, followed by a 2-electron oxidation to a quinone methide (BHTOH-QM). This highly electrophilic species alkylates cellular components such as proteins and possibly DNA. This molecular damage leads to activation of stress kinases (e.g., p38, JNK), and DNA damage response. If damage reaches a threshold level, caspase-3 is activated consistent with intrinsic apoptosis.

### Kinetic isotope labeling suggests t-butyl hydroxylation is rate limiting for BHT bioactivation

To identify which H-abstraction steps are rate limiting for BHT bioactivation, we exploited the primary kinetic isotope effect (KIE) used to study the enzymatic mechanisms of many CYP450 enzymes (44, 45). By substituting deuterium at the site of rate-limiting hydrogen abstraction, we hypothesized that BHT bioactivation may be sufficiently slowed to manifest as reduced toxicity. Unlabeled BHT was compared against two deuterated isotopologues: BHT-d3, bearing deuterium on the *p*-methyl group, and BHT-d21, deuterated at all positions except the *p*-methyl group (Figure 8C). As shown in Figure 8D, BHT-d21 showed approximately half the cytotoxicity of BHT-d0 at 20 and 40 µM doses. At 20 µM the KIE was not statistically significant (p = 0.23) likely due to the smaller overall difference and the assay being performed at a relatively early timepoint (5 h) prior to peak LDH release. However, at 40 µM the KIE of BHT-d21 reached statistical significance. Together with the structure-activity data showing that the tert-butyl substituent is required for toxicity, these results implicate hydrogen abstraction at the tert-butyl group as the initiating (and likely rate-limiting) step in CYP2B10-dependent BHT bioactivation.

## DISCUSSION

The BHT model represents an essential tool for investigating cellular mechanisms of lung epithelial repair and fibrosis. While multiple studies have suggested CYP450 activity is necessary for BHT’s pneumotoxicity (26, 46), here we show that AT1-derived CYP2B10 is capable of bioactivating BHT into a toxic QM. This leads to *in situ* protein alkylation, stress kinase activation, DNA damage response and intrinsic apoptosis in a cell autonomous manner. However, this newly discovered CYP2B10 biochemistry may have implications beyond murine lung injury models.

Firstly, relatively little is known about enzymatic oxidation of substituted phenols (e.g., *p*-cresols) into QMs. Perhaps the most studied example is raloxifene, which undergoes Cyp3a4-mediated activation into a QM followed by time-dependent inactivation of CYP3A4 through alkylation of a specific Cys residue (47, 48). Isotopic labeling with ^18^O2 and mutagenesis experiments supported a dehydrogenation reaction – rather than aryl epoxidation – en route to the raloxifene-derived QM that inhibits CYP3A4 (49, 50). Analogously, 4-hydroxylated metabolites of tamoxifen can be oxidized *in vitro* to a QM causing CYP450 alkylation and inhibition (51, 52). In contrast to QMs derived from other P450 enzymes, we did not identify BHT or BHTOH-modified peptides on CYP2B10 itself. Thus a natural resistance to alkylation may allow this enzyme to continue catalytic turnover without self-inactivation.

Secondly, features of CYP2B10-induced BHT bioactivation may facilitate future efforts at targeted cell ablation and bioremediation. For example, the CYP2B10-BHT pair may expand the toolbox for therapeutic suicide genes in allogeneic transplanted cells or chimeric antigen T-cells (CAR-T) as a method to arrest undesirable toxicities (53–55), particularly considering the lack of apparent bystander effect. With respect to bioremediation of BHT, CYP2B10 could potentially be utilized to chemically activate and thus covalently capture BHT out of a contaminated environment.

While SPAs such as BHT fall under the FDA health categorization of GRAS, our work demonstrates that at least one mammalian enzyme (CYP2B10) converts BHT into a highly toxic electrophilic species. Considering BHT biomagnification with contaminated aquatic environments and food supplies, our discovery provides a mechanistic pathway by which BHT could exhibit undesirable health effects. One relevant question is whether, do human CYP450s (including orthologue CYP2B6) exhibit similar reactivities towards BHT? If so, it seems feasible that BHT exposure could constitute a risk factor for chronic microalveolar injury that promotes ILDs such as idiopathic pulmonary fibrosis, a disease that is rarely explained by monogenic abnormalities. We hope that our discovery of enzymatic BHT bioactivation will pique the interest of the biochemical and environmental sciences communities, thus helping to address key outstanding questions and furthering the potential technologies relating to this unexpectedly novel biochemistry.

## LIMITATIONS OF THE STUDY

In the course of addressing a handful of hypotheses, our work raises many questions surrounding the interactions between p-cresols and CYP450 enzymes. For example, why does t-butyl hydroxylation appear necessary for bioactivation into an alkylating species? Drugs like bupropion undergo *t*-butyl hydroxylation by CYP2B enzymes (56), yet *t*-butyl hydroxylation is unlikely to be sufficient for complete bioactivation into a QM. Presumably BHTOH is the preferred substrate to undergo further oxidation by CYP2B10, although further work will be needed to address this outstanding issue.

Another key question is how BHTOH-QM triggers apoptosis? Is this through global stress or alkylation of a specific Cys site that triggers the apoptotic cascade? Two potential mitochondrial apoptotic triggers identified in our BHTOH site data include the resolving Cys230 of mitochondrial peroxiredoxin-3 (PRDX3) (57, 58) and Cys48 on voltage dependent anion channel-2 (VDAC2) (59). Both of these sites are considered ligandable in CysDB. It is conceivable that selective alkylation of PRDX3 and VDAC2 sensitizes mitochondria to endogenously produced reactive oxygen species, facilitating Bak-mediated mitochondrial permeabilization, cytochrome C release and intrinsic apoptosis as observed in our experiments.

## RESOURCE AVAILABILITY

For additional details or requests for resources/reagents, please contact the corresponding author. All unique/stable reagents generated in this study are available from the lead contact with a completed Materials Transfer Agreement.

## DATA AND CODE AVAILABILITY

Raw and processed data, and associated metadata, have been deposited to the ProteomeXchange consortium (PXD083806) via the MassIVE repository (ftp://massive-ftp.ucsd.edu/v14/MSV000103146/). All code used for data analyses is available at https://github.com/mt-forrester/11250_BHT_3T3_proteome. Any additional information required to reanalyze the data reported in this paper is available upon request.

## Supporting information

Supplementary Table S0 - README

## ACKNOWLEDGMENTS

This work was funded by the National Institutes of Health grants K08HL181195 (to MTF), K08CA263300 (to CEE), R01HL153375 and R01HL160939 (to PRT), Duke Department of Medicine Chairs Research Award (to MTF), Office of Physician Scientist Development Technician Support Award (to MTF) and Nanaline H. Duke Fund Strong Start Award (to MTF).

## AUTHOR CONTRIBUTIONS

MTF conceived the project, designed and executed experiments, analyzed data and wrote the manuscript. KMH and MWF performed experiments, analyzed data and edited the manuscript. CEE and PRT provided intellectual contributions and insights into project direction, analyzed data and edited the manuscript. NM, KT, RLF, JMM and AT provided intellectual contributions, insights into project direction and edited the manuscript.

