## Supplementary Information and Figures for "Alveolar Cytochrome P450 Mediates Butylated Hydroxytoluene-Induced Electrophilic Injury and Apoptosis"

**Supplementary Methods**

*Cloning and mutagenesis of Cyp2b10 and Cyp4b1* – Mouse lung was subjected to RNA isolation using the Qiagen RNAeasy kit followed by reverse transcription with Invitrogen SmartScript III using oligo-dT primers. The Cyp2b10 and Cyp4b1 gene products were subjected to restriction digestion (NheI / AgeI for Cyp2b10; XhoI / AgeI for Cyp4b1), ligated into pCMV-EGFP using T4 DNA ligase at 16C overnight with 4:1 molar ratio of insert:vector and transformed into NEB-Stable E coli (product number C3040H). Sequencing of colonies revealed ORFs matching to UniProt accessions Q9WUD0 and Q64462 for Cyp2b10 and Cyp4b1, respectively. These ORFs were transferred into an in-house optimized lentiviral vector (pFW series) consisting of a full EF1 promoter, multiple cloning site followed by T2A-hygR or T2A-EGFP for plasmids pFW2 and pFW3, respectively. All lentiviral plasmids were verified by whole plasmid sequencing. For mutagenesis of Cys436 to Ala, NEB basechanger protocol was employed with the indicated primers. All plasmids were verified by whole plasmid sequencing through Azenta / Genewiz.

*LC-MS/MS Sample Preparation* – Twenty µg of cell lysates were adjusted to 25 µL with 20% SDS in 50 mM TEAB buffer, pH 8.5, reduced by addition of 2.5 µL of 100 mM DTT and heating at 80 °C for 10 min. After cooling, samples were alkylated by addition of 2.5 µL of 250 mM iodoacetamide and incubation in the dark for 30 min. To each tube, 3 µL of 20% phosphoric acid was added followed by 200 µL of 90% MeOH/100 mM TEAB, and the samples were processed using S-trap micro devices (Protifi). Samples were digested with 1.5 µg of Sequencing Grade Modified Trypsin (Promega):protein for 1 h at 47 °C. Peptides were eluted with 40 µL TEAB, 40 µL 0.2% formic acid and 35 µL of 50% MeCN/0.2% FA. Lyophilized peptides were reconstituted in 80 µL of 0.1% formic acid. SPQC samples were prepared by mixing equal volumes of all samples. Finally, 3 µL of samples (including replicates of the SPQC) were loaded onto Evotips, followed by an addition of spiked with 50 fmol yeast ADH1 digestion (Waters Massprep).

*Liquid chromatography tandem mass spectrometry (LC-MS/MS) with data-independent acquisition* – Quantitative LC/MSMS was performed using an Evosep One LC coupled to a ThermoFisher Orbitrap Astral via a Nanospray Flex ionization source. The Evosep used a 30 sample-per-day method with a Pepsep 15 cm x 150 µm column (1.5 µm particle size) with a PepSep Sprayer and stainless steel (30 µm) emitter. The MS analysis used a 240,000 resolution Orbitrap precursor ion (MS1) scan from 380-1080 m/z, automatic gain control (AGC) target of 500% and maximum injection time (IT) of 50 ms, collected every 0.6 s in centroid mode. MS/MS was performed in the Astral analyzer using a DIA method with default charge state = 3, precursor mass range of 380-980 m/z,4 m/z isolation windows, AGC target of 500% and max IT of 6 ms. A normalized collision energy (NCE) of 28% for all MS2 scans, and a RF lens of 40% was used for MS1 and DIA scans. Individual samples were also analyzed using data-dependent acquisition (DDA). LC-MS/MS was performed as described above except that a 24,0000 resolution MS1 scan from 375-1500 m/z was collected every 1 s, followed by a ddMS2 Astral method with intensity threshold of 1E4, auto dynamic exclusion, 1.2 m/z isolation window, stepped collision energy of 24,26,28 %NCE, AGC target of 100%, maximum IT of 20 ms and first mass of 110 m/z.

*Analysis of DIA data* – Raw MS data was converted to *.htrms format using HTRMS converter and processed in Spectronaut 20.5 (Biognosys; 20.5.260227.92449). A spectral library was built using direct-DIA searches of all individual files (DDA + DIA) and used a Uniprot mouse database (UP000000589) downloaded on 04/08/2022 and appended with Cyp2b10 WT, Cyp2b10 C436A and as well as additional contaminant sequences using FragPipe (17,221 total entries). Search settings included trypsin/P specificity with up to 2 missed cleavages and peptide length from 7-52 amino acids with variable acetyl (protein N-term, carbamidomethyl(Cys) and BHT(OH)Cys modification. For DIA analysis, default extraction, calibration, identification, and protein inference settings were used. Data was filtered at a 1% precursor and protein group false discovery rate (q-value). Background imputation was optionally selected in Spectronaut. Local normalization of quantified precursors,[1] and protein roll-up with the MaxLFQ algorithm[2] were used within Spectronaut. Diagnostic ions were also extracted from DIA data using the "--extract 217.16, 235.17" option in DIA-NN [3]. Tab-delimited XIC files containing retention time, DIA isolation window boundaries, and per-scan intensities for each diagnostic ion were read into R (v4.x). Raw scan-level intensities were organized into DIA cycles by detecting resets in the isolation window lower boundary (Window.Low). Consecutive subcycles were then paired into full DIA cycles, and the diagnostic-ion intensities within each full cycle were summed to produce a single aggregated intensity value per cycle, with the corresponding retention time taken as the mean of the constituent scan times. For visualization, XICs for m/z 235.17 and 217.16 were filtered to retention times ≥ 20 minutes. Vertically offset chromatograms (for each replicate) were displayed in a faceted grid using ggplot2. For quantitation, intensities were binned to a common 0.05-minute retention time grid by rounding each cycle's mean RT to the nearest 0.05 min and averaging intensities falling within the same bin. Per-condition summary statistics (mean ± SD across three replicates) were then computed at each binned time point. Both individual-replicate and condition-summary data were exported as CSV files.

*Analysis of DDA data* – DDA raw files were searched in FragPipe v24.0 (MSFragger v4.4.1) against the mouse SwissProt proteome with appended common contaminants and reversed decoys (2026-03-16-decoys-reviewed-contam-UP000000589.fas; decoy prefix rev_). Two complementary searches are outlined below.

*Open search (global modification profiling)* – A mass-tolerant "open" search was performed in FragPipe v24.0 (MSFragger) using the default "Open" workflow against the same database. The precursor mass window was -150 to +500 Da with delta-mass localization enabled; precursor true tolerance and fragment tolerance were 20 ppm with mass calibration/optimization. Cleavage was fully tryptic (stricttrypsin, K/R) with up to 2 missed cleavages; peptides 7-50 residues, 500-5000 Da. Carbamidomethylation of cysteine (+57.02146) was fixed; oxidation of methionine (+15.9949) and protein N-terminal acetylation (+42.0106) were variable (<=3 per peptide), with N-terminal methionine clipping allowed. Open-search artifacts were addressed with Crystal-C, PSMs were validated with PeptideProphet using the extended mass model, and results were filtered to 1% FDR in Philosopher. The global modification landscape was profiled with PTM-Shepherd (default parameters), yielding per-modification PSM summaries.

*BHTOH labile/offset search (diagnostic-ion-enforced site identification)* – The BHT-quinone-methide Michael adduct (BHTOH, +234.16199 Da) was searched as a labile mass offset (labile_search_mode = labile; mass_offsets = 0.0/234.16199) restricted to cysteine (restrict_deltamass_to = C), with delta-mass fragment localization enabled (localize_delta_mass = 1; labile b/y ion series) and the delta mass reported as a variable modification. Spectra were required to contain BHT-QM diagnostic oxonium evidence: fragments m/z 217.159240 and 235.169805 at a summed relative intensity >=10% of the base peak (diagnostic_intensity_filter = 0.1). Cleavage was fully tryptic (stricttrypsin, K/R) with up to 1 missed cleavage; peptides 7-50 residues, 500-5000 Da. Precursor and fragment tolerances were +-20 ppm with mass calibration/optimization (calibrate_mass = 2), isotope-error search 0/1/2, and deisotoping enabled. No fixed modifications and no additional variable modifications were applied (carbamidomethylation was not modeled); N-terminal methionine clipping was allowed and up to 3 variable modifications per peptide were permitted. A match required >=4 matched fragment ions and >=2 sequence-specific ions. PSMs were validated with PeptideProphet and filtered to 1% protein-level FDR in Philosopher (--sequential --prot 0.01; PSM/peptide 1%); MSBooster and Percolator were disabled. Adduct sites were localized with PTMProphet (FRAGPPMTOL = 10 ppm, MINPROB = 0.5). Label-free site quantification used IonQuant with MaxLFQ (m/z tolerance 10 ppm, RT tolerance 0.4 min, ion-level FDR 1%, >=1 ion and >=3 scans per feature, minimum localization probability 0.75, normalization on, match-between-runs off).

*Gene set enrichment analysis from DIA data* – Pre-ranked GSEA was performed on the WT+BHT vs WT+Vehicle proteome contrast using clusterProfiler (v4.20.0). Proteins were ranked by a signed significance metric, sign(log2FC) × −log10(p-value), from the limma result; gene symbols were mapped to Entrez IDs (org.Mm.eg.db v3.23.0), collapsed to one entry per gene then sorted in decreasing order. Mouse MSigDB Hallmark gene sets (collection "H") were retrieved with msigdbr (v26.1.0) and supplied as a TERM2GENE table. Enrichment was computed with clusterProfiler::GSEA() (fgsea pre-ranked algorithm) using gene-set size limits of 15–500, 0.05 significance cutoff, Benjamini–Hochberg adjustment, eps = 0, and a fixed random seed (set.seed(42)).

### ***Primers used for cloning***

| Gene | Template | Destination | Forward Primer | Reverse Primer |
|---|---|---|---|---|
| Cyp2b10 | Mouse lung cDNA (oligo dT) | pCMV-EGFP NheI / AgeI | ATTAGCTAGCACCATGGAGCCCAGTG | AATAACCGGTCGGGCCAAGAAGCAGATC |
| Cyp4b1 | Mouse lung cDNA (oligo dT) | pCMV-EGFP XhoI / AgeI | TATACTCGAGACCATGGCGCTCAGC | TTATACCGGTTTTCCAGACCCAGGG |
| Cyp2b10 | pCMV-2b10-EGFP | pFW2/3 XbaI / EcoRI | TTAATCTAGAGCCACCATGGAGCCC | AATTGAATTCTCGGGCCAAGAAGCAG |
| Cyp4b1 | pCMV-4b1-EGFP | pFW2/3 XbaI / MfeI | ACTATCTAGAGCCACCATGGCGC | TAATCAATTGTTTTCCAGACCCAGGGCC |

| | | | | |
|---|---|---|---|---|
| Cyp2b10 | pFW2/3-Cyp2b10 | C436A SDM | AAAGCGCATTGCGCTTGGTGAAAGC | CCTGTTGAGAAGGGC |

### *Key Resources Table*

| Reagent | Manufacturer | Product # | Usage Notes (if applicable) |
|---|---|---|---|
| Butylated hydroxytoluene (BHT) | Sigma | W218405 | |
| 2-tert-Butyl-4-methylphenol | Sigma | B97208 | |
| 2-tert-Butyl-6-methyl-phenol | Sigma | B97607 | |
| 2,4-Dimethylphenol | Thermo | 408450050 | |
| 2,6-Di-tert-butyl-4-ethylphenol | Sigma / Ambeed | AMBH93E4C47F | |
| Butylated hydroxytoluene-d3 | HY-Y0172S2 | | |
| Butylated hydroxytoluene-d21 | MedChemExpress | HY-Y0172S | |
| Polybrene Solution (10 mg/mL) | Sigma | TR-1003 | |
| Thiazolyl Blue Tetrazolium Bromide (MTT) | Sigma | M2128 | |
| Goat anti-Mouse IgG2A Alexa 647 | Invitrogen | A21241 | 1:500 in 2% BSA/TBST, secondary for anti-Cyp2b immunostaining in IgG2A-deficient mice (e.g., C57BL/6) |
| Donkey anti-Rabbit IgG Alexa 488 | Invitrogen | A21206 | 1:500 in 2% BSA/TBST |
| Donkey anti-Rabbit IgG Alexa 594 | Invitrogen | A32754 | 1:500 in 2% BSA/TBST |
| Anti-Cyp2b Mouse IgG2a | Origene | TA504328S | WB 1:1000 in 5% milk/TBST, IF 1:1000 with MeOH fixative or PFA + antigen retrieval in citrate buffer pH 6.0 |
| Cleaved caspase 3 mAb | CST | 9664 | IF 1:500, PFA fixation, antigen retrieval not necessary |
| Hopx rabbit polyclonal Ab | Invitrogen | PA5-90538 | IF 1:500, PFA fixation, antigen retrieval |
| LEL 488 - Lycopersicon Esculentum (Tomato) Lectin DyLight 488 | Invitrogen | L32470 | Final 10 µg/mL in PBST; used after secondary Ab staining for 10-20 min |
| GAPDH mouse mAb | Proteintech | 60004 | WB 1:2000 in 5% milk / TBST |
| p38 mAb | CST | 9212 | WB 1:1000 in 5% milk / TBST |
| cJun mAb | CST | 9165 | WB 1:1000 in 5% milk / TBST |
| SAPK/JNK mAb | CST | 9252 | WB 1:1000 in 5% milk / TBST |
| Chk1 mAb | CST | 2360 | WB 1:1000 in 5% milk / TBST |
| PARP mAb | CST | 9542 | WB 1:1000 in 5% milk / TBST |
| phospho p38 (Thr180/Tyr182) mAb | CST | 4511 | WB 1:1000 in 2% BSA / TBST |
| phospho JNK (Thr183/Tyr185) mAb | CST | 4671 | WB 1:1000 in 2% BSA / TBST |
| phospho cJun (Ser73) mAb | CST | 9164 | WB 1:1000 in 2% BSA / TBST |
| yH2AX mAb | CST | 9718 | WB 1:2000 in 2% BSA / TBST |
| phospho Chk1 (Ser 345) mAb | CST | 2348 | WB 1:1000 in 2% BSA / TBST |
| phospho-RPA32 (Ser33) mAb | CST | 10148 | WB 1:1000 in 2% BSA / TBST |
| Sequencing Grade Modified Trypsin | Promega | V5111 | |
| CytoTox 96 LDH Assay | Promega | G1780 | |
| Caspase Glo 3/7 Assay | Promega | G8091 | |

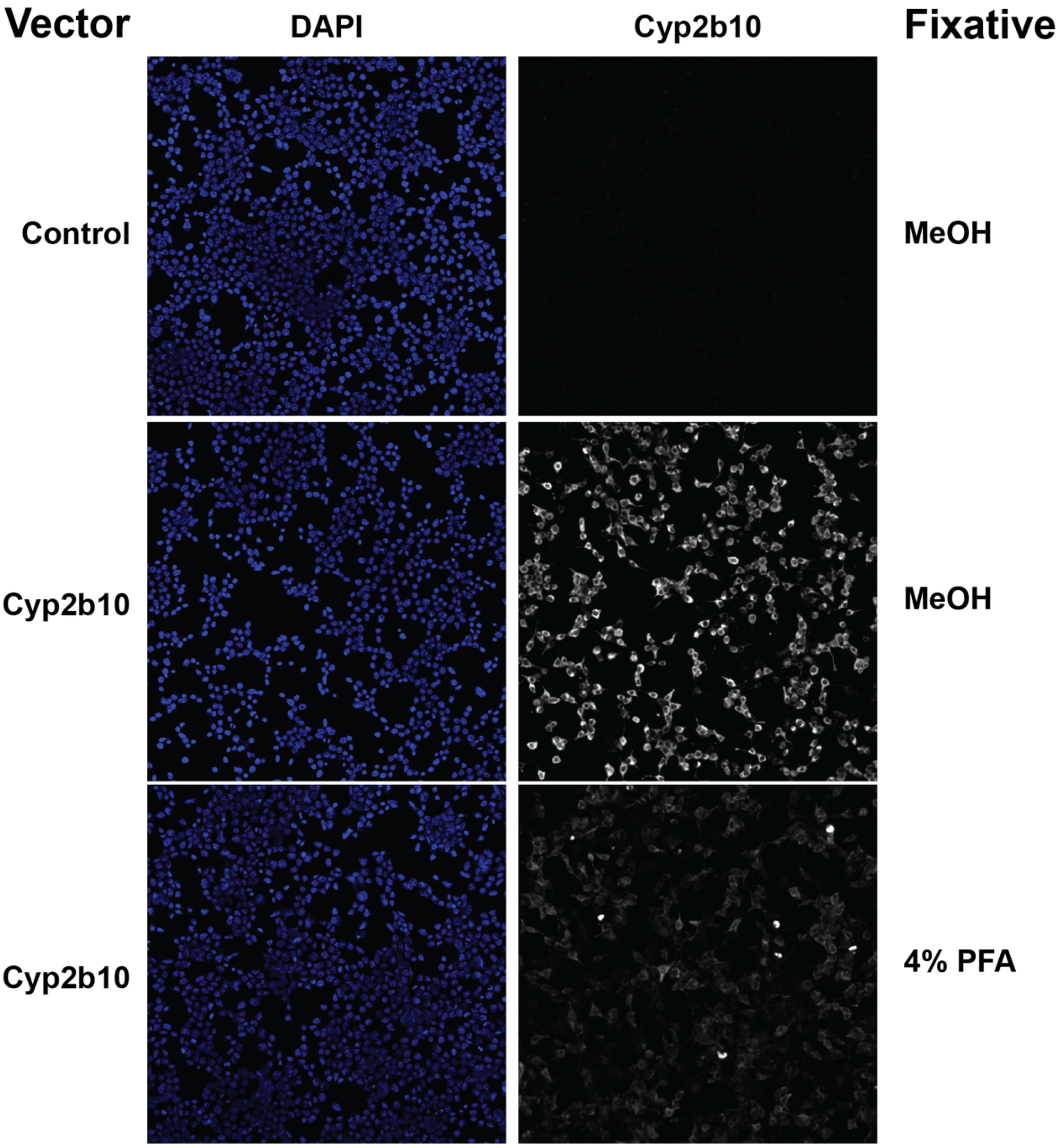


**Figure S1. Validation of commercially available antibody against Cyp2b (Origene mouse monoclonal IgG2A)**. HEK293T cells were seeded onto poly-lysine coated coverslips, then transiently transfected with plasmids encoding either EGFP or Cyp2b10-T2A-EGFP. Twenty-four hours later, cells were washed with PBS and fixed in either cold MeOH or 4% PFA for 5 minutes. Cells were permeabilized with tris-buffered saline containing 0.1% Triton X-100 for 5 min at RT, blocked in TBST + 2% BSA, and stained with anti-Cyp2b10 IgG2A antibody at 1:1000 dilution overnight. Secondary goat anti-mouse IgG2A Alexa-Fluor 647 Ab was used at 1:1000 dilution. Coverslips were mounted and imaged by confocal microscopy at 20x magnification. Expression of EGFP was assessed to confirm transfection. Shown are DAPI (nuclei) and Cyp2b10 immunoreactivity. Results are representative of n = 3 separate experiments.

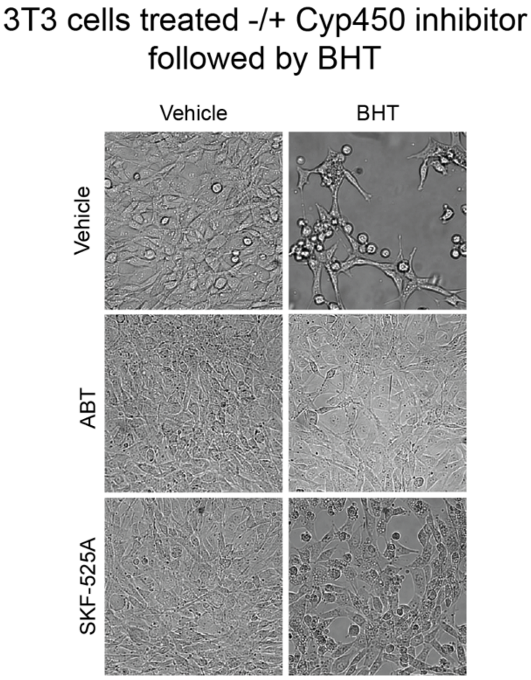


**Figure S2. Brightfield microscopy of NIH-3T3 cells treated with Cyp450 inhibitors followed by BHT.** Prior to harvesting, cells were imaged by brightfield microscopy at 20x magnification. Shown is a single representative replicate from data in Figure 3B.

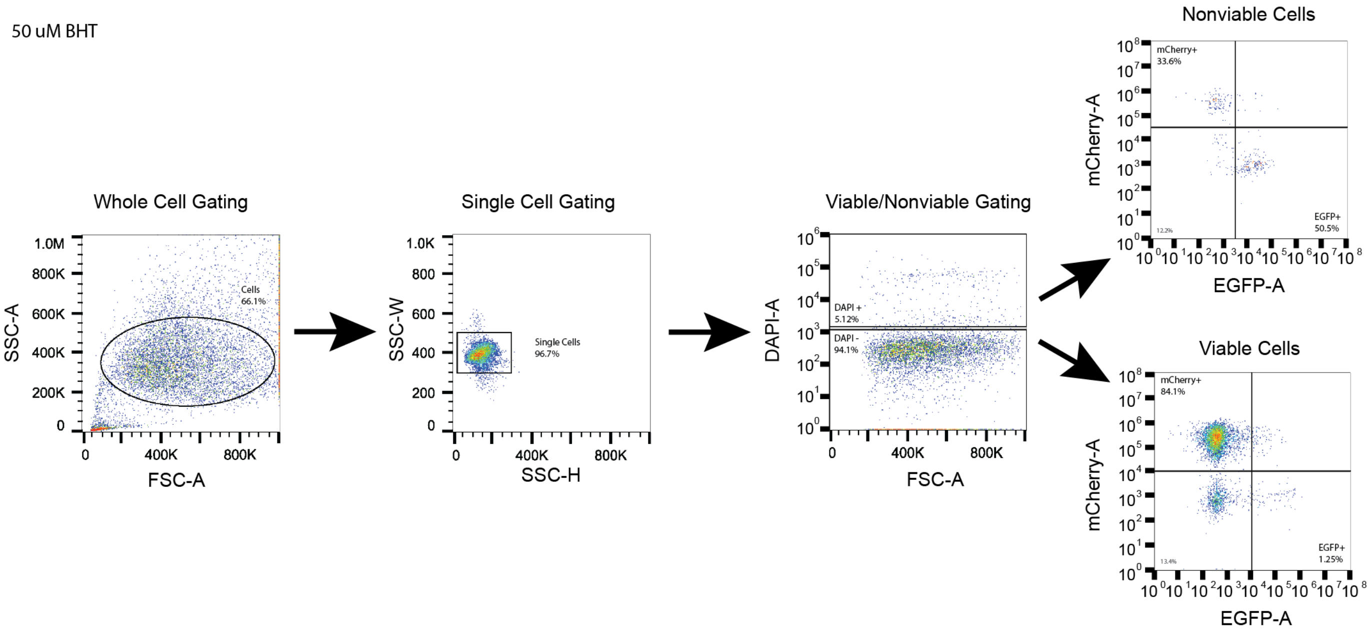


**Figure S3. Sample gating strategy for bystander flow cytometry experiment.** Shown is an individual replicate of the sample treated with 50 μM BHT.
